# Binned Relative Environmental Change Indicator (BRECI): A tool to communicate the nature of differences between environmental niche model outputs

**DOI:** 10.1101/672618

**Authors:** Peter D. Wilson

## Abstract

Niche models are now widely used in many branches of the biological sciences and are often used to contrast the distribution of favourable environments between regions or under changes in environmental conditions such as climate change. Evaluating model performance and selecting optimal models is now accepted as best-practice, and a number of methods are available assist this process. One aspect of ENM application which has not received as much attention is developing methods to communicate the degree and nature of changes between model outputs (typically as raster maps). The method described in this paper, Binned Relative Environmental Change Index (BRECI), seeks to address this shortfall in communicating model results.

---

Models of the realised niche of organisms are now routinely used to relate species occurrence to environment in fields as diverse as ecology (Ranjitkar et al. 2014, Baker et al. 2021), population genetics (Bariotakis et al. 2016, Dellicour et al. 2017), biogeography and comparative studies (Smith and Donoghue 2010, Alvarado-Serrano and Knowles 2014), and ethnobotany (Gaikwad et al. 2011). Just as diverse is the terminology for these models which includes ecological or environmental niche model (ENM, a term used in this paper), species distribution model (SDM), resource selection models (RSMs) and habitat suitability models (HSMs).

ENMs are frequently used to contrast environmental suitability between two time periods to assess the potential impact of changed environmental conditions, particularly those related to climate change. Comparison of ENM output maps for the two times may provide insight into the degree of change, and its spatial distribution, and thus inform management responses. However, few studies have provided a way of communicating the nature of differences between ENM outputs. Given their widespread use and importance to conservation assessment and planning, it is vital that those specialists creating ENMs assist decision-makers and stakeholders by clearly communicating the outputs of sometimes complex ecological models (Stillman et al. 2016, Sofaer et al. 2019, Baker et al. 2021).

In this note I introduce a new graphical tool, the Binned Relative Environmental Change Indicator (BRECI), to assist in understanding and communicating the nature of differences between pairs of ENM output maps. I define the method for computing BRECI, illustrate its application, briefly discuss its use in relation to evaluating model performance and statistical map comparison, and provide *R*-code for its computation.

## A recipe for BRECI

When two rasters are available for comparison and both have the same raster geometry (grid origin, grid cell size, map projection and coordinate system) and the same environmental suitability scale, proportional change values may be computed for an arbitrary set of environmental suitability bins which partition the suitability scale into values from “low” to “high”. Both the number of bins used and the suitability values defining bin widths are arbitrary parameters which may be adjusted to suit the task at hand. When these parameters are set, computing a suite of BRECI values proceeds as follows:

1. For each of two ENM output rasters to be compared, *Map1* and *Map2*, replace cell values with the index of the bin into which they fall;
2. Generate a cross-tabulation between grid cell values
3. For each bin, *i*, compute the proportional or relative change in the number of grid cells between the two maps relative to the number cells in that bin on *Map1*:

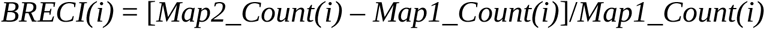

*or*
4. For each bin, *i*, compute the proportional or relative change in the number of grid cells between the two maps relative to the number cells in raster:

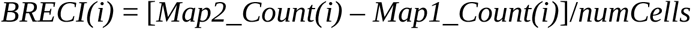

Normalising the score in each bin by the number of calls in that category in Map1 (see Step 3) exaggerates scores and scores in each bin can be greater than +/- 1. This may be useful when assessing relative scores between bins when bins have similar scores. Selecting to normalise all bins by the number of cells (Step 4) in the raster ensures that scores in each bin range between −1 and +1.

The sign of the *BRECI* value in each bin indicates gain (positive values) or loss (negative values) of environmental suitability. The majority of ENM tools produce suitability scores transformed to a 0 (no suitability) to 1 (maximum suitability) scale, or may be easily scaled to this range. For output scaled in this way, a reasonable but arbitrary number of bins is five equal-sized bins using bin cut-points at 0.2, 0.4, 0.6, 0.8. The bins can be loosely but reasonably interpreted as shown in Table 1. A similar scheme can be applied to maps based on any suitability scale.

**Table 1:** Suggested default bin ranges and an heuristic interpretation of suitability values included within each bin. The function BRECIplot in the R-package BRECI allows users to supply their own subdivision of suitability values with associated interpretive labels

| **Bin** | **Range** | **Interpretation** |
| --- | --- | --- |
| 1 | 0 – 0.2 | Very low suitability |
| 2 | 0.2 – 0.4 | Low suitability |
| 3 | 0.4 – 0.6 | Moderate suitability |
| 4 | 0.6 – 0.8 | High suitability |
| 5 | 0.8 – 1.0 | Very high suitability |

An R-package, *BRECI*, is available in GitHub to produce BRECI plots (https://github.com/peterbat1/BRECI).

## Application of the method

The use of BRECI plots is illustrated in Figure 1 by applying it to three synthetic pairs of rasters representing “before” and “after” ENM outputs covering three scenarios: overall loss, little change and overall gain of suitability. The application of the BRECI method to these three examples illustrates how a BRECI plot can highlight the nature of predicted environmental changes.

**Figure 1:**
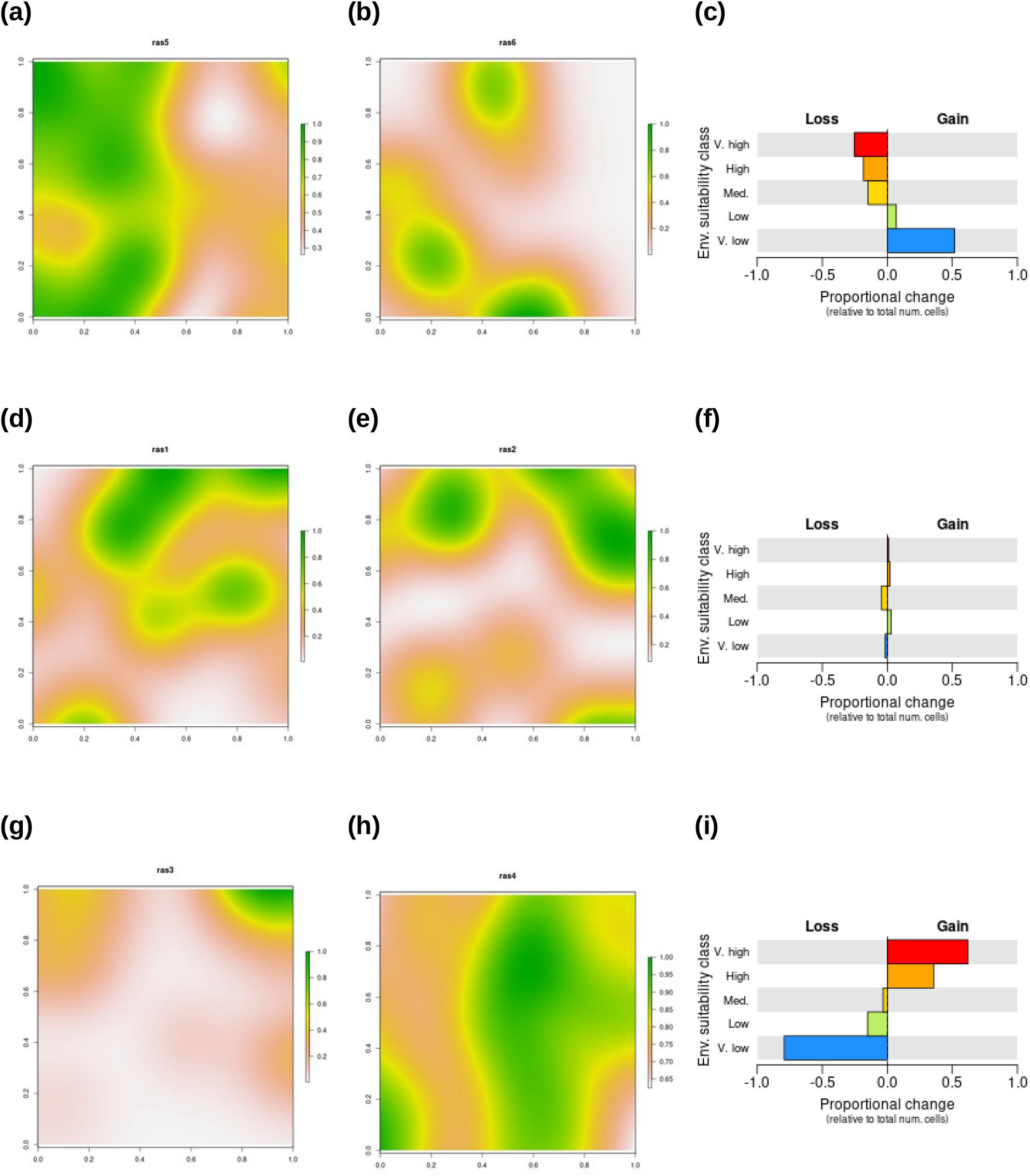
Example applications of the BRECI method to three synthetic ENM map pairs. (a), (b) and (c). “Before” and “After” rasters and BRECI plot showing substantial loss of environmental suitability. (d), (e) and (f). “Before” and “After” rasters and BRECI plot showing little change in environmental suitability. (g), (h) and (i). “Before” and “After” rasters and BRECI plot showing substantial gain in environmental suitability.

A BRECI plot shows an overall trend in suitability scores between two raster. It is ‘aspatial’: it does not show where on the map transitions have occurred. That could be very important information, but it becomes a real challenge to make those assessments without assistance. The function *gainLossMaps* in the BRECI R-package can provide the needed spatial insights. The function produces an overall view of areas of gains and losses as separate overall gain and overall loss maps. It also produces a set of gain or loss pairwise transition maps for each possible transition between pairs of bins. The use of this function is illustrated in Figure 2.

**Figure 2:**
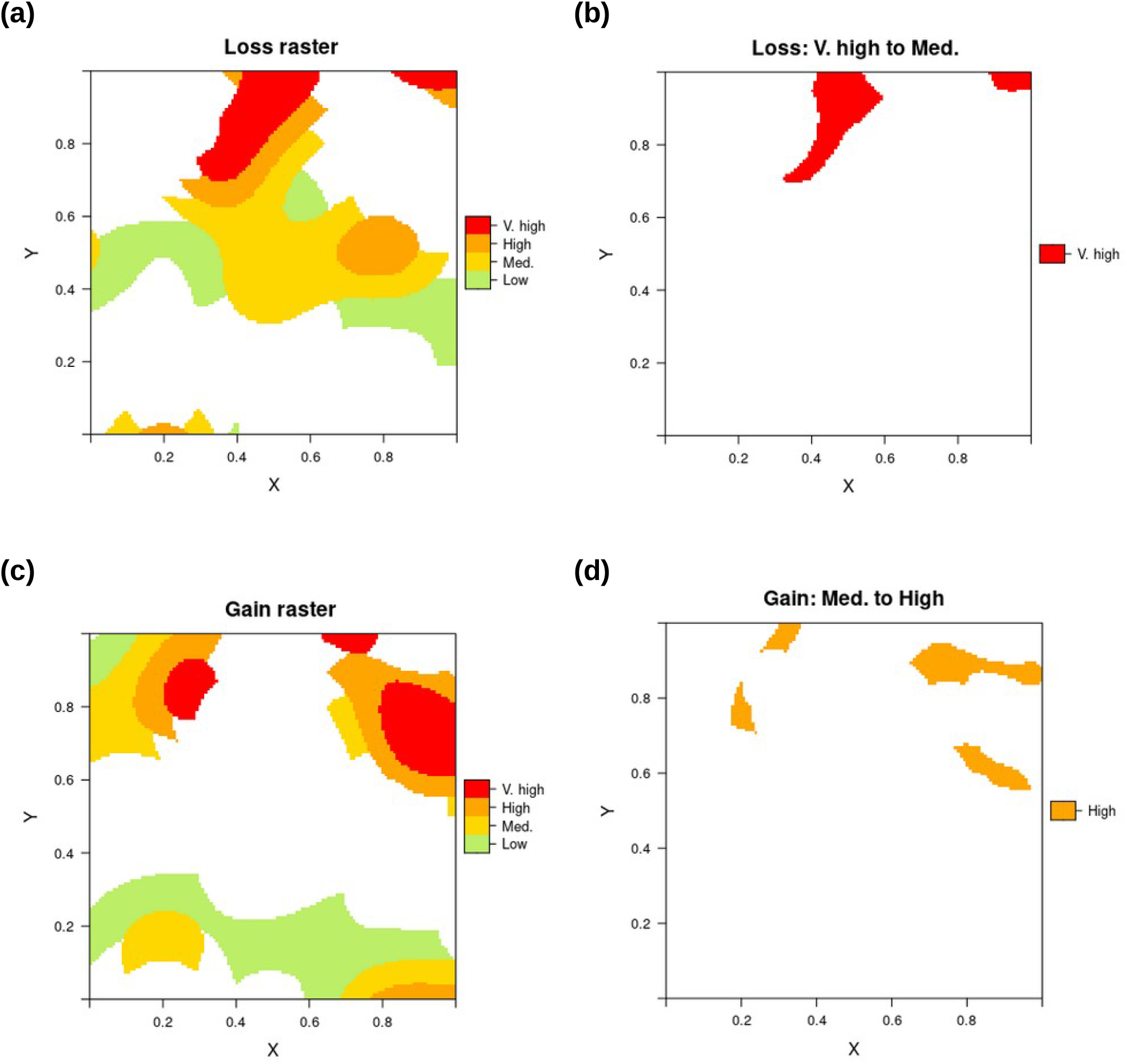
Example outputs from the function gainLossMaps in the R-package BRECI. (a) Transition map showing regions which lost suitability. (b) Areas in the overall loss map shown in (a) which transitioned from Very High suitability to Medium suitability. (c) Transition map showing regions which gained suitability. (d) Areas in the overall gain map shown in (c) which transitioned from Medium to High suitability. The number of pairwise contrasts grows rapidly with the number of bins chosen to compute the BRECI plot.

## Other methods of map comparison

Communicating complex and often subtle information presented in maps is challenging (Board and Taylor 1977). Responding to this challenge, practitioners in the spatial sciences have developed many methods for map comparison (Boots and Csillag 2006, Foody 2006, 2007, Jones et al. 2016). Methods for comparing raster maps, including ENM outputs, may be grouped according to whether they used binary rasters, or directly compare the continuous-valued rasters output by ENM software.

Converting ENM output to binary form requires the application of a threshold value. Selecting an appropriate threshold is a significant challenge (Liu et al. 2005, Freeman and Moisen 2008, Nenzén and Araújo 2011) and may mask important differences in the distribution of marginal environmental suitability. Even though it is problematic, thresholding is widely used and several methods are available to compare binary maps. Simple counts of pixels classed as “suitable” or counts of pixels considered suitable in both maps may be made (e.g. Mellick et al. 2014). Alternatively, more sophisticated statistical comparisons are also possible (Boots and Csillag 2006, Robertson et al. 2007).

Methods for the direct un-thresholded comparison of raster maps such as ENM output maps include statistical tests of pixel (grid cell) differences (Levine et al. 2009) and other moments of pixel values (Wealands et al. 2005, Jones et al. 2016), measures of niche overlap and adaptations of the Hellinger distance between probability distributions (Warren et al. 2010, Wilson 2011), threshold-free measures of range shift and overlap (Kou et al. 2014), and simple correlation between rasters (Wealands et al. 2005, Syphard and Franklin 2009)

One motivation for direct spatially-referenced comparison of model output rasters is that model fits for the same taxon may produce the same or very similar performance metrics yet display different spatial distributions of suitability scores (Wilson 2011). An additional motivation is that it is frequently not valid to use statistical measures of model performance to compare models for different taxa (Guisan et al. 2017) or between models (Hand 2009, Parker 2011) but the projection of models into geographical space does permit similarities and differences to be examined across taxa and between model types. A further motivation is to represent the nature of differences between maps in an objective and simple to guide decision making.

The Binned Relative Environmental Change Indicator (BRECI) is designed to provide a simple but objective representation of map differences. It allows two aspects of the difference between two maps of environmental suitability: (a) overall visualisation of the magnitude and direction of change, and (b) representation of the nature of that change including the spatial distribution of regions of gain or loss. In addition, BRECI plots may be produced for the whole model extent or any arbitrary region within it. The most important feature of this method is that it avoids the use of binary thresholds which have the potential to give a misleading indication of the spatial extent of, and changes in, suitability.

Overall visualisation of ENM map differences can be reduced to a statement about “gain” or “loss” of environmental suitability. Often such statements are made by applying a threshold to the continuous-valued maps produced by ENMs, turning them into binary maps. BRECI allows users to easily understand the degree of “gain” or “loss” across a spectrum of environmental suitability classes and so avoid information loss associated with simple binary thresholds. This makes it possible to provide a ranking of species into categories such as “severe loss”, “moderate loss”, etc. In many instances this is acceptable for triage decisions or for indicating the way members of species ensembles or ecological assemblages may face disparate futures. It is also possible to produce a BRECI plot for the entire extent of a fitted model, or arbitrarily defined spatial sub-sets, allowing assessments of range-wide change to be contrasted with, say, changes within a particular management region or jurisdiction.

The second aspect, visualising the nature of differences, is achieved by partitioning values in the map into bins or environmental suitability classes. Clearly communicating complex information in map form can be challenging, requiring trade-offs and choice between many different elements of map design having regard to the constraints of human visual perception (Board and Taylor 1977). The relative gain or loss in each bin allows users to see the way suitability changes across the spectrum of values present in the two maps. For example, it is possible to discriminate between pair-wise comparisons showing major changes in all bins with those showing large overall change but where change is dominated by differences in a particular environmental suitability class. These insights into the nature of changes in suitability could then be included in management decisions: Should greater weight be given to taxa suffer large losses of high suitability versus those predicted to face losses of areas of marginal suitability?

## Conclusions

Communicating the implications of outputs from complex ecological models is critical to achieve good conservation outcomes, but can be challenging and requires additional tools to translate and communicate model outputs (Stillman et al. 2016). ENMs have become an important tool in impact assessments and conservation planning (Sofaer et al. 2019, Baker et al. 2021). A tool like BRECI can play a useful role in communicating the magnitude and nature of changes in ENM outputs.

BRECI has a number of strengths including: ease of computation, an intuitive interpretation, the ability to convey aspects of difference not easily represented in statistically orientated measures, avoids the potential for errors introduced by applying a threshold to make binary maps, and it may be applied to the output of *any* modelling method which produces environmental suitability scores on raster maps. Naturally, there are some limitations which include: it is not a substitute for model performance measures, it uses an arbitrary sub-division into suitability classes or bins, and it can only compare maps with values on the same suitability scale (e.g. MaxEnt logistic scale versus MaxEnt logistic scale) and the same raster geometry (e.g. matching grid cell size, grid origin, map projection and coordinate system). Experience has shown that placing a BRECI plot alongside the two maps being compared is a powerful heuristic tool for communicating the nature of differences between the maps.

## Acknowledgements

The first version of BRECI was developed while I was undertaking post-doctoral work at the Macquarie University, Sydney, Australia under the guidance of Prof. Lesley Hughes and Prof. Michelle Leishman. An early version was applied to assessing the relative impact of climate change on over 2,000 weed species in Australia (www.peterwilson.id.au). The current version was developed to support the Restore & Renew project at the Royal Botanic Gardens, Sydney (www.restore-and-renew.org.au) lead by Dr Maurizio Rossetto.

## References

Alvarado-Serrano, D. F., and L. L. Knowles. 2014. Ecological niche models in phylogeographic studies: applications, advances and precautions. Molecular Ecology Resources 14:233–248.

Baker, D. J., I. M. D. Maclean, M. Goodall, and K. J. Gaston. 2021. Species distribution modelling is needed to support ecological impact assessments. Journal of Applied Ecology 58:21–26.

Bariotakis, M., K. Koutroumpa, R. Karousou, and S. A. Pirintsos. 2016. Environmental (in)dependence of a hybrid zone: Insights from molecular markers and ecological niche modeling in a hybrid zone of Origanum (Lamiaceae) on the island of Crete. Ecology and Evolution 6:8727– 8739.

Board, C., and R. M. Taylor. 1977. Perception and Maps: Human Factors in Map Design and Interpretation. Transactions of the Institute of British Geographers 2:19.

Boots, B., and F. Csillag. 2006. Categorical maps, comparisons, and confidence. Journal of Geographical Systems 8:109–118.

Dellicour, S., C. Kastally, S. Varela, D. Michez, P. Rasmont, P. Mardulyn, and T. Lecocq. 2017. Ecological niche modelling and coalescent simulations to explore the recent geographical range history of five widespread bumblebee species in Europe. Journal of Biogeography 44:39–50.

Foody, G. M. 2006. What is the difference between two maps? A remote senser’s view. Journal of Geographical Systems 8:119–130.

Foody, G. M. 2007. Map comparison in GIS. Progress in Physical Geography 31:439–445.

Freeman, E. A., and G. G. Moisen. 2008. A comparison of the performance of threshold criteria for binary classification in terms of predicted prevalence and kappa. Ecological Modelling 217:48–58.

Gaikwad, J., P. D. Wilson, and S. Ranganathan. 2011. Ecological niche modeling of customary medicinal plant species used by Australian Aborigines to identify species-rich and culturally valuable areas for conservation. Ecological Modelling 222:3437–3443.

Guisan, A., W. Thuiller, and N. E. Zimmermann. 2017. Habitat Suitability and Distribution Models. Cambridge University Press, Cambridge, UK.

Hand, D. J. 2009. Measuring Classifier Performance: A coherent alternative to the Area Under the ROC Curve. Machine Learning 77:103–123.

Jones, E. L., L. Rendell, E. Pirotta, and J. A. Long. 2016. Novel application of a quantitative spatial comparison tool to species distribution data. Ecological Indicators 70:67–76.

Kou, X., Q. Li, C. Beierkuhnlein, Y. Zhao, and S. Liu. 2014. A new tool for exploring climate change induced range shifts of conifer species in China. PLoS ONE 9:e98643.

Levine, R. S., K. L. Yorita, M. C. Walsh, and M. G. Reynolds. 2009. A method for statistically comparing spatial distribution maps. International Journal of Health Geographics 8:7.

Liu, C., P. M. Berry, T. P. Dawson, and R. G. Pearson. 2005. Selecting thresholds of occurrence in the prediction of species distributions. Ecography 28:385–393.

Mellick, R., P. D. Wilson, and M. Rossetto. 2014. Demographic history and niche conservatism of tropical rainforest trees separated along an altitudinal gradient of a biogeographic barrier. Australian Journal of Botany 62:438–450.

Nenzén, H. K., and M.B. Araújo. 2011. Choice of threshold alters projections of species range shifts under climate change. Ecological Modelling 222:3346–3354.

Parker, C. 2011. An analysis of performance measures for binary classifiers. Pages 517–526 2013 IEEE 13th International Conference on Data Mining. IEEE Computer Society, Los Alamitos, CA, USA.

Ranjitkar, S., R. Kindt, N. M. Sujakhu, R. Hart, W. Guo, X. Yang, K. K. Shrestha, J. Xu, and E. Luedeling. 2014. Separation of the bioclimatic spaces of Himalayan tree rhododendron species predicted by ensemble suitability models. Global Ecology and Conservation 1:2–12.

Robertson, C., T. A. Nelson, B. Boots, and M. A. Wulder. 2007. STAMP: spatial–temporal analysis of moving polygons. Journal of Geographical Systems 9:207–227.

Smith, S. A., and M. J. Donoghue. 2010. Combining historical biogeography with niche modeling in the Caprifolium Clade of Lonicera (Caprifoliaceae, Dipsacales). Systematic Biology 59:322–341.

Sofaer, H. R., C. S. Jarnevich, I. S. Pearse, R. L. Smyth, S. Auer, G. L. Cook, T. C. Edwards, G. F. Guala, T. G. Howard, J. T. Morisette, and H. Hamilton. 2019. Development and delivery of species distribution models to inform decision-making. BioScience 69:544–557.

Stillman, R. A., K. A. Wood, and J. D. Goss-Custard. 2016. Deriving simple predictions from complex models to support environmental decision-making. Ecological Modelling 326:134–141.

Syphard, A. D., and J. Franklin. 2009. Differences in spatial predictions among species distribution modeling methods vary with species traits and environmental predictors. Ecography 32:907–918.

Warren, D. L., R. E. Glor, and M. Turelli. 2010. ENMTools: a toolbox for comparative studies of environmental niche models. Ecography 33:607–611.

Wealands, S. R., R. B. Grayson, and J. P. Walker. 2005. Quantitative comparison of spatial fields for hydrological model assessment––some promising approaches. Advances in Water Resources 28:15– 32.

Wilson, P. D. 2011. Distance-based methods for the analysis of maps produced by species distribution models. Methods in Ecology and Evolution 2:623–633.

